# Social closeness modulates brain dynamics during distrust anticipation

**DOI:** 10.1101/2022.03.08.483524

**Authors:** Said Jiménez, Roberto Mercadillo-Caballero, Diego Angeles-Valdes, Juan José Sánchez-Sosa, Jairo Muñoz-Delgado, Eduardo A. Garza-Villarreal

## Abstract

Anticipation of trust from a partner with high social closeness generates the preconditions for cooperative and reciprocal interactions to occur. However, if there is uncertainty in the interaction because the partner is a stranger or because the person has mistrusted us on another occasion, an aversive experience is generated. Hence, low social closeness or mistrust makes people keep track of the other’s behavior. In case of experiencing deviations from social norms, the person will monitor the intentions of the partner and update their priors regarding their social preferences. The anterior insula (AIns) seems to be sensitive to these social norm violations in functional magnetic resonance imaging (fMRI) experiments, however, the monitoring of partners with different levels of social closeness has not been investigated. In our study we wanted to find the brain regions related to (dis)trust anticipation from partners who differ in their level of social closeness. For this, we designed an experiment in which participants played an economic decision game with three people (trustors): A computer, a stranger, and a real friend. We covertly manipulated their decisions in the game so that they unexpectedly distrusted our participants. Using task-fMRI, our whole-brain analysis found that the AIns was active during the anticipation of the decisions from human partners (humans vs computer), but not during anticipation between high and low social closeness (friend vs stranger). However, using a psychophysiological interaction analysis, we found increases in functional coupling between the AIns and regions in the “mentalizing” network (such as temporal regions and parieto-occipital cortices) during trust anticipation between a high versus low social closeness partner. These results suggest that there may be a modulation of the AIns activity, specifically for high social closeness trustors, by regions that encode the intentions underlying the truster behavior.

## 1. Introduction

The anticipation of trust from an individual towards a person with high social closeness (i.e. a close friend) could be considered the status quo of interactions between members of the same group (Fehr & Schurtenberger, 2018; Krueger et al., 2020). From borrowing a pen from a friend to requesting a friend to endorse a bank loan, these are situations which reflect trust that is anticipated between members of the same social network. The expectation of trust from a person who shares high social closeness with another, allows their behavior to be predicted very accurately and generates the preconditions for cooperative and reciprocal interactions to occur (Fareri, 2019; Krueger et al., 2020). However, if the other person is a stranger (someone with low social closeness) or if the context could imply risk due to social uncertainty, it is appropriate to anticipate that trust may not occur by default. Thus, anyone anticipating the trust of a partner needs to adapt their expectations and prepare from possible deviations from the social norm of trust, depending on information such as social closeness (Krueger & Hoffman, 2016).

The trust placed from one person to another is regarded as a type of social reward (Izuma et al., 2008). The anticipation of this affiliative behavior seems to involve the activation of brain regions such as the ventral striatum (VS), which has been implicated in reward processing and has shown significant differential activity when people trust in-group versus out-group members (Hughes et al., 2017). During the anticipation of a reinforcer, the Salience Network (Schneider et al., 2018; Wilson et al., 2018) (SN), anchored in the anterior cingulate cortex (ACC) and the anterior insula (AIns), seems to be recruited to orchestrate motivational and attentional processes. In the context of a potentially prosocial interaction, the SN may help detect compliance or violation of social norms and could guide the decision to respond with reciprocity or else, to update the initial belief regarding the social preferences of the other person (Krueger & Meyer-Lindenberg, 2019).

Economic games are one of the main tools to study prosocial behavior and the associated neural circuits. In particular, the Trust Game allows exploring the brain mechanisms that underlie both the truster’s ability to place trust in others and the trustee’s decision to respond reciprocally (Rilling & Sanfey, 2011). Although there are numerous studies that have investigated brain circuits involved in the decisions of both roles (truster and trustee), little is known regarding the influence of social closeness on trust anticipation of the trustee. Even though it is true that the effect of social inputs has been found mainly related to the activity of isolated brain regions during cooperation, competition or approval (e.g., such as the VS or medial prefrontal cortex (mPFC)) (Fareri et al., 2015; Fareri & Delgado, 2014a, 2014b; Sip et al., 2015), the neural dynamics during trust anticipation from a partner with high social closeness (a friend) compared to one with low closeness (stranger), is currently an open question.

In this study, we aimed to explore the neural dynamics of the AIns during trust anticipation from a friend compared to a stranger, as well as to evaluate whether the response of brain regions involved in anticipation is modulated by the partner’s social closeness. We are particularly interested in the activity of the right AIns, due to its role in the expectation of possible social norm violations, and its direct influence on regions that coordinate the executive and affective processes that underlie reciprocity or defection (Bellucci et al., 2018; Pärnamets et al., 2020). For this purpose, we manipulated social closeness in a Trust Game by introducing three partners: a computer (non-social control), a stranger (out-group member), and a friend considered close (ingroup member). In the trustee’s role, the subjects independently anticipated the trust from their peers, and subsequently decided whether to reciprocate or not. While in the trustor’s role, the partners decided whether to invest a monetary amount in the trustee, which could then result in higher profits for both. To ensure that our participants experienced the violation of the social norm and the associated emotional uncertainty, we covertly manipulated the behavior of the three trustors so that they randomly decided to distrust the participant in some of the trials.

Due to the AIns’ role in the interoceptive signals representation and its belonging to the Salience Network (Critchley, 2008), our exploratory hypothesis was that, given the uncertainty regarding the behavior of its peers, the subjects will show a more aversive experience depending on the social closeness of the trustor, particularly, for the peer of greater social closeness, reflected in the AIns bold signal increase. Likewise, presumably due to the need to update the initial beliefs about the social preferences of their peers, we hypothesized that the AIns will be functionally coupled with limbic cortices and with mentalizing regions such as angular gyrus and posterior cingulate cortex in the default mode network (Krueger & Meyer-Lindenberg, 2019). According to the neuropsychological framework of third-party punishment, the neural dynamics, would allow the individual to provide an estimate of the severity of the latent damage caused by a possible transgression, as well as infer the trustor’s motives for mistrust (Krueger & Hoffman, 2016).

## 2. Material and methods

### 2.1. Participants

We recruited 30 healthy subjects (15 female), all reported being right-handed, and ranged between 19 to 33 years old. No subject disclosed a neurological history or psychiatric illness. The participants attended our study with a “close” friend (n = 30), who had the following characteristics: 1) they were match-gender friends paired with the participant, 2) they were not a relative, and 3) they were not a romantic or sexual partner. Exclusion criteria were related to MRI safety such as claustrophobia or ferromagnetic metals in the body. The study was approved by the ethics committee of the Instituto Nacional de Psiquiatría Ramón de la Fuente Muñiz in Mexico City. All participants and their friends gave written consent for the study, and we followed the guidelines of the Declaration of Helsinki.

### 2.2. Procedure

A researcher explained participants they would be scanned with MRI while playing an economic game (experimental fMRI task) with three partners: 1) their friend, 2) stranger (same sex) and 3) a computer. They were told that their friend was going to play with them in an isolated room, and that the stranger was another person who already knew the game and was waiting in another room for the moment, although they never met the stranger. We also told them that the computer partner was programmed to make decisions that benefit it. In reality, the participants were deceived as they did not play with anyone, and the responses and behavior of the friend, stranger, and computer were all programmed a priori to control the response variability. This deception was necessary as an experimental manipulation to ensure the effect of social closeness not to be affected by the real-time responses, to induce a level of distrust in the participant, as well as to control the timing of the study. To ensure the level of social closeness that the participants and their friends reported, the two responded to the IOS scale “Inclusion of the Other in Self” (Aron et al., 1992) without observing their friend’s responses. It consists of seven pairs of circles that vary in the degree of overlapping and represent the subjective social closeness that one individual perceives with respect to another. Highly overlapping circles suggest high social closeness, while distant circles indicate the opposite. The participants were asked to answer the IOS scale about the friends, the stranger, and the computer. After the verbal explanation of the economic game task, the participants were trained first outside the MRI scanner, then inside to get accustomed. Following training, the experiment began and lasted for 1 hour. At the end of the experiment, the participants and their friends were told about the deception. All of our participants were given the opportunity to be eliminated from the study if they did not agree with any of the manipulations and deceptions performed by the investigator, however, none of our subjects chose that option.

### 2.3. Experimental task

The task was programmed in PsychoPy 1.84.2 (Peirce, 2008) and the participant observed the task on a viewer adapted for use inside the scanner, and responded by pressing two buttons on the response pad Lumina PAIR Pad of Cedrus, one of the buttons was used to the decision to “to pay back”, while the other was to “not to pay back”. The participants played the role of trustee in a Trust Game (Berg, 1995) (the task) against three trustors of different degrees of social closeness: computer (control), stranger (low), and friend (high). Each trial included 4 phases: 1) promises, 2) trust anticipation, 3) decision, and 4) feedback. During the promise phase, participants had to promise their partners how often they would reciprocate; during the trust anticipation phase, participants had to wait for their partners’ decision about giving them money. In the decision phase, the participant had to decide whether or not to pay back to his trustor and, finally, during the feedback phase, payments for the trial were indicated depending on the decisions of the participants or the trustor’s response (Figure 1). The game consisted of 24 trials (8 for each partner) using hypothetical rewards: the trustor (computer, stranger, or friend) expressed his trust by investing $2 (Mexican pesos) in the trustee (participant), the trustee anticipated their partner’s decision for 6 seconds; if the trustor trusted, the $2 would be multiplied by 5 and delivered to the trustee, while the trustor ran out of money. Later, if there was an investment, the trustee had to decide whether to return half to the trustor (trustee $5, trustor $5) or keep all the money (trustee $10, trustor $0). If there was no investment, the trustee received nothing in that trial and waited for her next partner (trustee $0, trustor $2). The three trustors (friend, stranger, and computer) were presented in a pseudorandom order and their decisions were programmed to randomly trust 6 out of 8 trials and distrust 2 out of 8.

**Figure 1.**
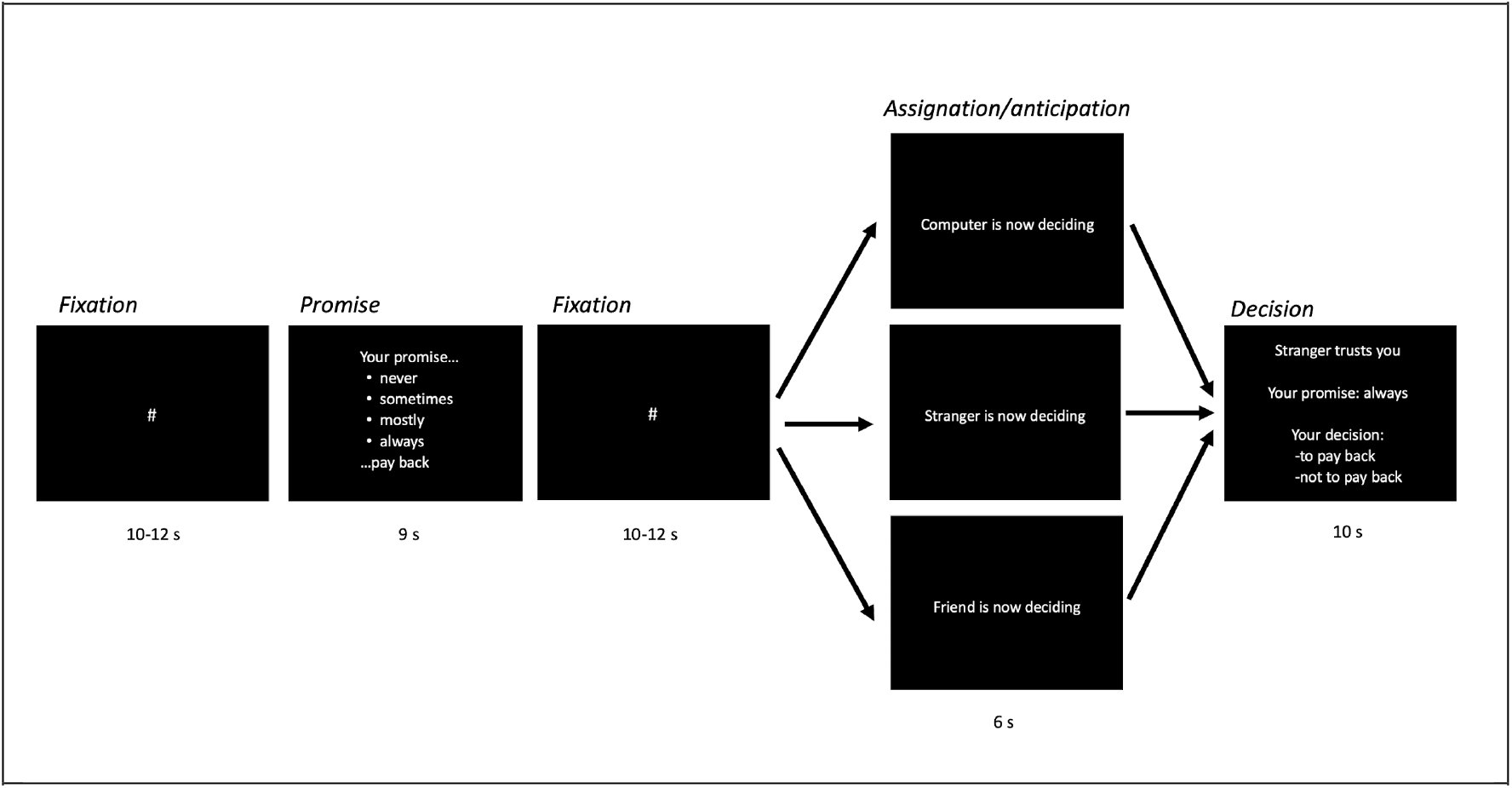
Task structure and timeline. From left to right the order and duration of the phases of a typical trial are shown; first, a fixation point was shown, then participants could make a promise regarding their payment frequency (Promise phase). After another fixation point, the subjects waited for 6s their investor’s decision, who could be: the computer, the stranger, or the friend (Anticipation phase). In order for the participants to experience distrust of the investor, the decisions of the three trustors were programmed to randomly invest in 6 out of 8 trials in the participant, and in 2 out of 8 not to invest. During the decision phase, the participants decided whether to pay or keep the money.

### 2.4. Image acquisition and preprocessing

Brain images were acquired using a Phillips Ingenia 3T MR system scanner (Philips Healthcare, Best, The Netherlands, and Boston, MA, USA), with a 32 channel dS Head coil. Functional data were acquired using a T2*-weighted echo-planar image sequence, with a repetition time (TR)/echo time (TE) = 20000/30 ms, flip-angle = 75°, and inversion recovery for cerebrospinal fluid suppression. A total of 510 axial slices were acquired with an isotropic resolution of 3 mm, field of view = 240 mm, and acquisition matrix = 80 x 80. The structural data were acquired by means of a T1-weighted sequence with TR/TE = 7/3.5 ms, flip angle = 8°, field of view = 240 x 240, 1.0 mm isotropic voxels, acquisition plane = sagittal.

### 2.5 MRI analysis

MRI data were analyzed using FSL 6.0.1 (FMRIB’s Software Library). For preprocessing, each 4D volume was motion and slice timing corrected, and normalized onto MNI common brain space (Montreal Neurological Institute, EPI Template, voxel size 2 mm x 2 mm x 2 mm). Data were then smoothed using a Gaussian filter (full width half maximum = 6 mm) and highpass filtered with sigma = 50(s). In order to examine the effect of social closeness on the BOLD signal of the brain regions involved in the trust anticipation, a first-level GLM was performed with 12 regressors, 6 movement regressors, 3 regressors of interest modeled the signal during the anticipation phase, and they lasted 6s each. One regressor was included to model the anticipation of the computer’s decision, a second regressor for the stranger, and a third regressor for the friend. Then, 3 regressors of no interest were included during the promises phase (9s), during the decision phase (10s), and during the promise phase control condition, which showed for 9s the message that said: “you can play without promises”. As part of the first-level analysis, three contrasts based on the hypothesis of interest were estimated: 1) the difference during the anticipation of the stranger investor compared to the computer (Stranger > Computer), 2) the difference during the anticipation of the friend investor compared to the computer (Friend > Computer), and 3) the difference during trust anticipation from a high compared to a low social closeness partner (Friend > Stranger). Finally, for statistical inference, we performed a one-sample t-test for each contrast as a higher-level analysis, the normalized statistical images were thresholded using clusters determined by Z>2.3 and a (corrected) cluster significance threshold of p=0.05 (Worsley, 2001).

### 2.6. Psycho-Physiological Interaction (PPI) analysis

To investigate the neural dynamics during trust anticipation, two PPI analyses were performed. We investigated if the functional connectivity of two ROIs of interest, the right AIns, and Parietal Inferior, was greater during the anticipation of a friend’s decision compared to the anticipation of a stranger. Very close to these areas, activation has been reported in tasks involving anticipation or prediction of prosocial behavior (Baumgartner et al., 2009; Morelli et al., 2014), as well as in the processing of physiological and psychosocial stressors (Kogler et al., 2015). The analyses were performed in FMRIB-FSL using the time series of the right AIns and Parietal Inferior activity as seed ROIs for each participant (O’Reilly et al., 2012), then a PPI model was fitted with 7 regressors as a first-level analysis. The first was the task regressor (PSY) that included the anticipation phase for the stranger and friend’s decisions, it had a duration of 6s, a weight of −1 was included for stranger’s anticipation and 1 for friend’s anticipation so that this regressor embodied the contrast Friend > Stranger; the second regressor was the physiological (PHY), for this, the right AIns and Parietal Inferior time-series activity during the entire task were used. The third regressor was the interaction between the psychological and physiological regressors (PSY*PHY). The remaining 4 regressors of the PPI model were covariates of no interest, three of them modeled the other task’s phases, one for the promises phase (9s), another for the control condition without promises (9s), and another for the anticipation of the computer’s decision (6s). The 4th of the no-interest regressor was used to model the shared variance between the anticipation phase of the friend’s decision and that of the stranger, it had a duration of 6s and included a weight of 1 for the anticipation of the two investors. To identify group-level activations, we performed a one-sample t-test as a higher-level analysis. The normalized statistical images were thresholded non-parametrically using the same parameters referred to in the whole-brain analysis.

### 2.7. Behavioral data analysis

Effects of promises and social closeness on the decision to reciprocate with trustors (pay back) were evaluated using a multilevel model with a binomial error distribution (Gelman & Hill, 2006), which was programmed in R with the lme4 package (Bates et al., 2015). To model the decision to pay back, the next categorical predictors were included as population-level effects (fixed effects): promises (with two levels: without promises/with promises), social closeness (with three levels: computer/stranger/friend), and the interaction promises by social closeness. The post hoc differences between trustors were analyzed using p-values adjusted with Tukey correction. The model also included the effect of social closeness at the individual level (random effects), to consider within-subject variability.

## 3. Results

### 3.1. Social closeness

To verify the assumption that the MRI participant experienced a similar degree of social closeness to their friends, we related the responses that the MRI participant and their friends gave to the IOS scale. We found a positive significant correlation between the subject and friend subjective levels of social closeness (r = 0.438, 95% CI: 0.09 - 0.69). Therefore, social closeness between the MRI participants and their friends was met.

### 3.2. Behavioral results

In the Trust Game, the participants had high levels of reciprocity as they decided to pay back to the trustors in 72% of trials regardless of social closeness and promises. However, the multilevel model and post hoc comparisons showed significant effects of social closeness in the reciprocity decision: the percentage that participants decided to pay back was statistically higher for the friend than the computer and stranger (Figure 2B). A significant increase in the decision to pay back was also found when participants made a promise compared to when they did not (Figure 2A). There were no significant interaction effects between promises and social closeness.

**Figure 2.**
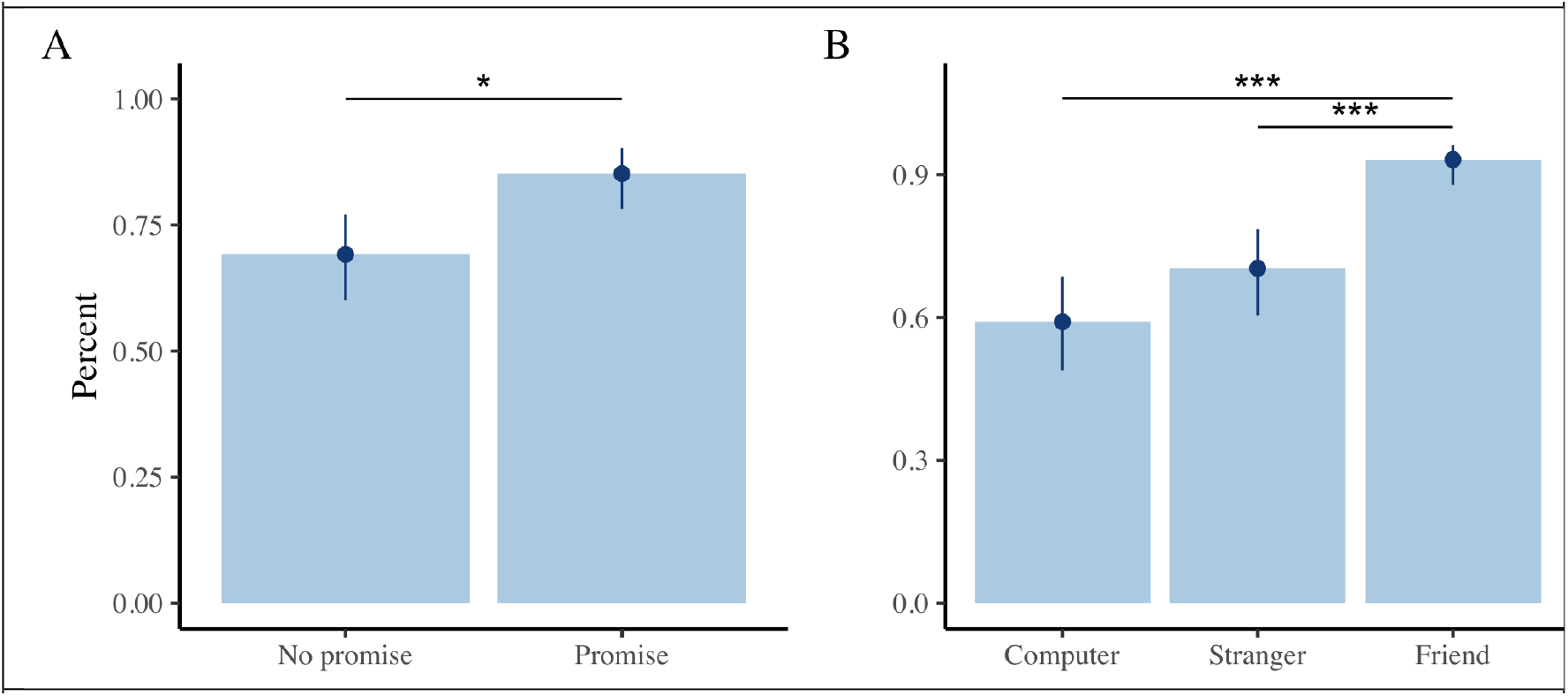
Effects of social closeness on reciprocity expressed by the decision to pay back in the Trust Game. A) Percentage of times that MRI participants decided to pay back when they made a promise compared to when they did not. B) Percentage of payments depending on the level of social closeness of the truster, the friend was paid significantly more than the stranger and the computer. *** p < 0.001.

### 3.3. fMRI main effects

The whole-brain analysis identified the brain regions that showed significant activation during the anticipation of the truster’s decisions, the contrast Computer-Stranger recruited activation of the basal ganglia: caudate nucleus and putamen (Figure 3A). The difference in activity during anticipation of the friend’s decision compared to the computer’s decision, expressed by the contrast Computer-Friend, showed the maximum activation peak in the Lingual gyrus, also it revealed significantly activated clusters that included the angular gyrus, cuneus, precuneus, putamen and the AIns (Figure 3B). Finally, the contrast Stranger-Friend found activation in the supplementary motor area (SMA), the middle frontal gyrus (MFG), the parietal lobe, and the angular gyrus (Figure 3C). Coordinates of the peak activations for all contrasts are shown in Table 1.

**Figure 3.**
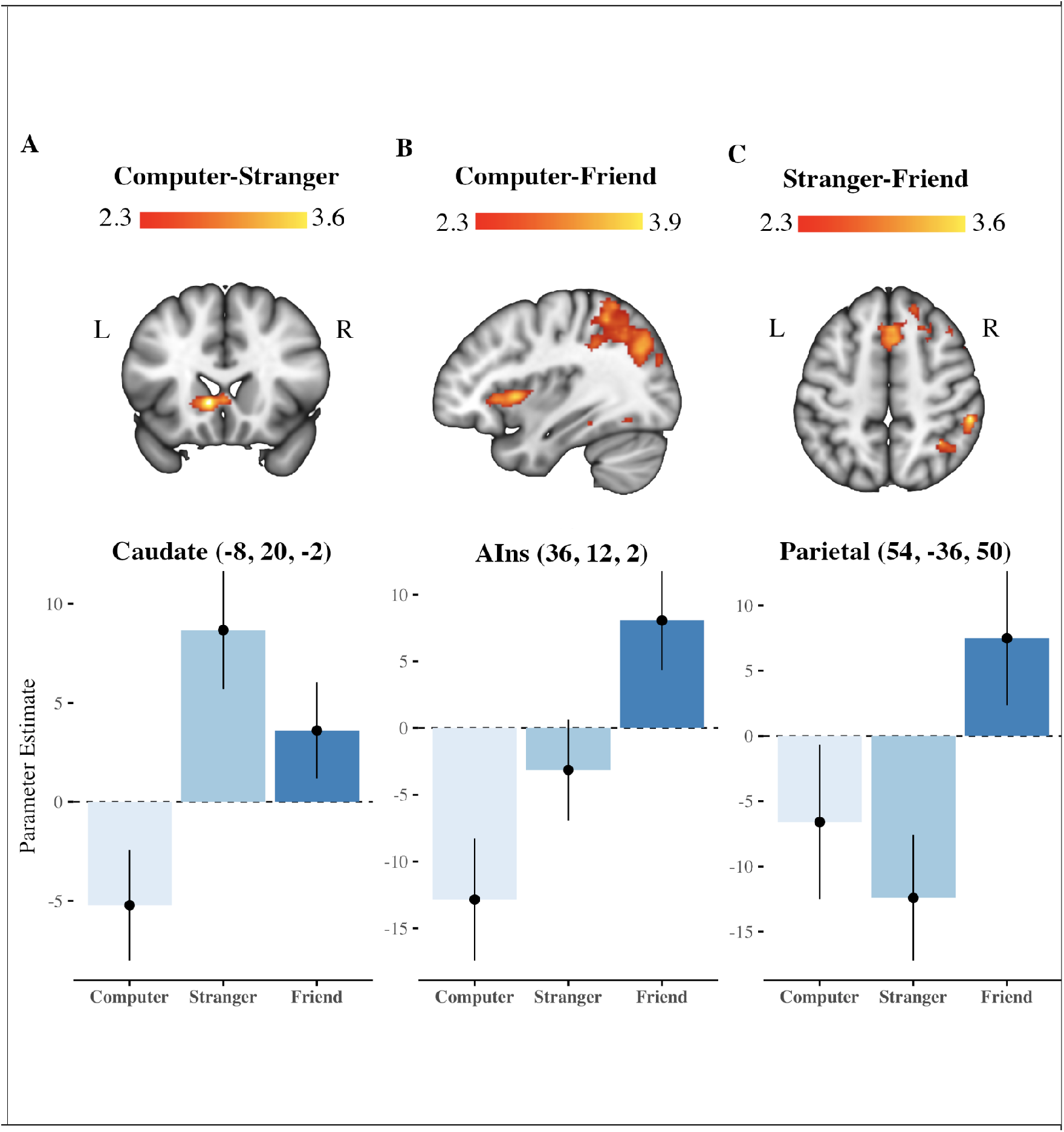
Neural regions involved in trust anticipation depending on the trustor’s social closeness Whole-brain analysis regions involved during trust anticipation were detected with the 3 contrasts of interest. A top. Coronal view of the statistical map for the Computer-Stranger contrast, the voxels with higher activation during the trust anticipation phase are represented on an intensity scale between 2.3 < z < 3.6. A bottom. This shows the value of the parameter estimated at the peak of activation depending on the social closeness. B top. Sagittal view of the statistical activation map for the Computer-Friend contrast, the voxels with the highest activation (highest Z-score) during the anticipation phase of trust are represented on an intensity scale between 2.3 < z < 3.9. B bottom. it represents the value of the parameter estimated at the peak of activation depending on the trustors’ social closeness. C top. Axial view of the activation statistical map for the Stranger-Friend contrast during the trust anticipation phase, plotted on an intensity scale between 2.3 < z < 3.6. C bottom. Representation of the estimated parameter in the maximum activation depending on social closeness.

**Table 1.** Whole-brain main effects

| Contrast | Region | BA | Coordinates (mm) |  |  | Z-score |
| --- | --- | --- | --- | --- | --- | --- |
|  |  |  | x | y | z |  |
| Stranger > Computer | Caudate | 48 | -8 | 20 | -2 | 3.68 |
|  | Putamen | 49 | -20 | 8 | -12 | 3.63 |
|  | Caudate | 48 | 0 | 16 | 0 | 3.56 |
|  | Putamen | 49 | -32 | 0 | 0 | 3.08 |
|  | Putamen | 49 | -24 | 2 | 4 | 2.97 |
|  | Putamen | 49 | -18 | 12 | 2 | 2.97 |
| Friend > Computer | Angular gyrus | 7 | 28 | -50 | 44 | 3.88 |
|  | Lingual gyrus | 19 | -20 | -54 | -10 | 4.00 |
|  | Cuneus | 18 | -12 | -86 | 26 | 3.44 |
|  | Precuneus | 7 | -12 | -62 | 66 | 3.41 |
|  | Anterior Insula | 13 | 36 | 12 | 2 | 3.65 |
|  | Putamen | 49 | -32 | -8 | -4 | 3.39 |
| Friend > Stranger | Supplementary motor area | 6 | 6 | 22 | 64 | 3.60 |
|  | Supplementary motor area | 8 | 4 | 14 | 48 | 3.45 |
|  | Supplementary motor area | 8 | 2 | 22 | 48 | 3.38 |
|  | Middle frontal gyrus | 8 | 42 | 22 | 46 | 3.32 |
|  | Supplementary motor area | 6 | 16 | 14 | 64 | 3.28 |
|  | Middle frontal gyrus | 8 | 26 | 24 | 44 | 3.25 |
|  | Parietal inferior | 40 | 54 | -36 | 50 | 3.60 |
|  | Parietal inferior | 7 | 34 | -52 | 50 | 3.60 |
|  | Middle occipital gyrus | 7 | 32 | -72 | 40 | 3.46 |
|  | Angular gyrus | 39 | 46 | -58 | 44 | 3.34 |
|  | Parietal inferior | 39 | 36 | -50 | 42 | 3.10 |
|  | Parietal superior | 39 | 40 | -58 | 58 | 3.08 |

### 3.4. PPI Results

Finally, the investigation of brain dynamics with the PPI model revealed a significant interaction between the anticipation of a trustor’s high vs. low social closeness and the time series of the right AIns. Specifically, during trust anticipation of the friend compared to the stranger, significantly greater functional coupling was found between the right AIns and the Fusiform Gyrus (FG). Likewise, the cluster of significant functional connectivity with the AIns, extended through the Angular Gyrus (AG) and Middle Occipital Gyrus (Figure 4A). Similarly, the right inferior parietal increased its functional connectivity with the FG, and the inferior/middle temporal gyrus, during trust anticipation of a friend versus a stranger (Figure 4B and Table 2).

**Figure 4.**
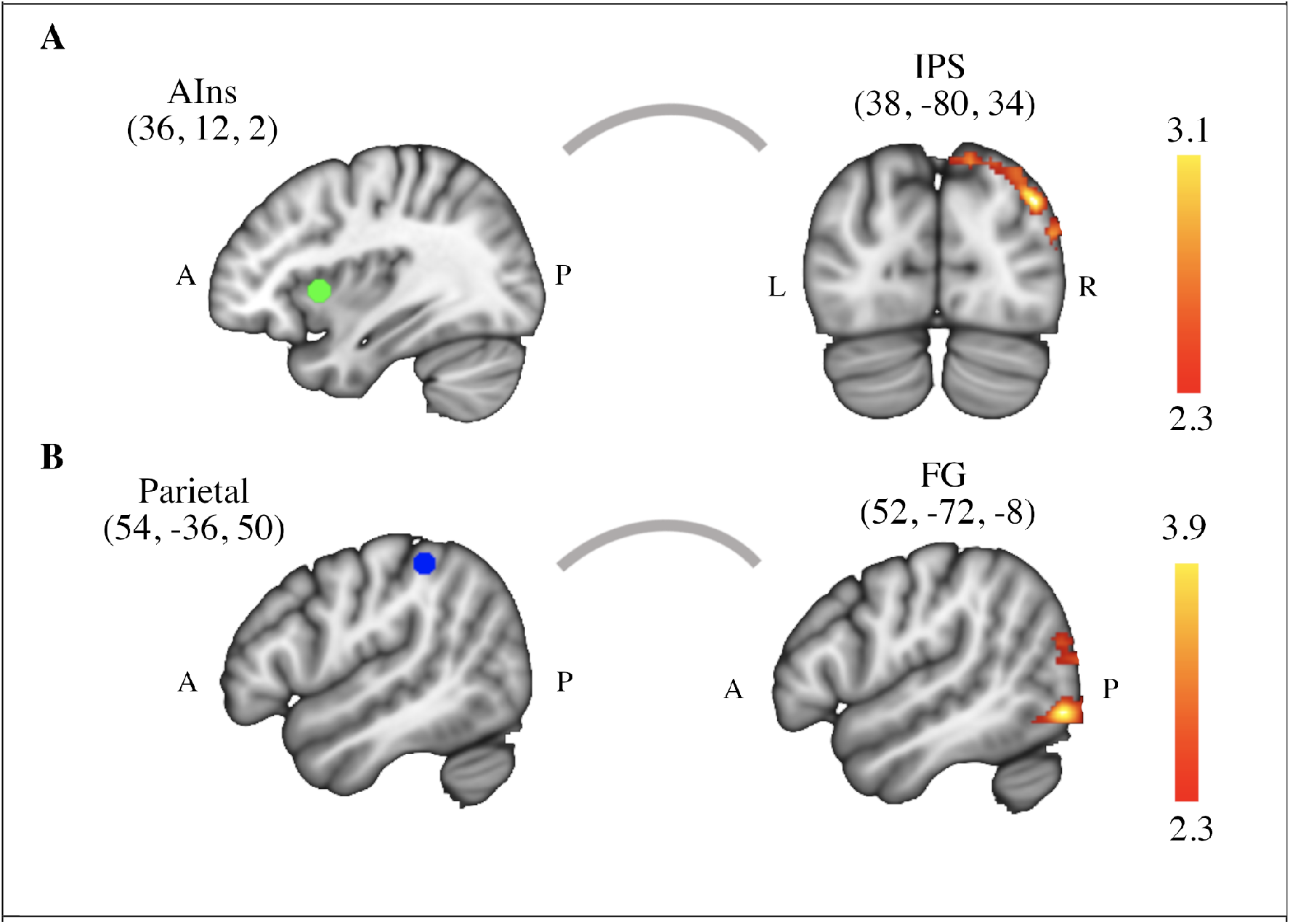
Psychophysiological interaction (PPI) results. A) The right AIns (AIns) was the seed region (MNI coordinates 36, 12, 2). Greater functional connectivity was found between AIns and IPS during anticipation of the friend’s decision compared to the stranger. B) The other seed was the right parietal cortex (MNI coordinates 54, −36, 50) which was coupled with the FG and the inferior/middle temporal gyrus during trust anticipation of a friend versus a stranger. Non-parametrically thresholded images using clusters determined by Z> 3.1 and a (corrected) cluster significance threshold of p = 0.05.

**Table 2.**
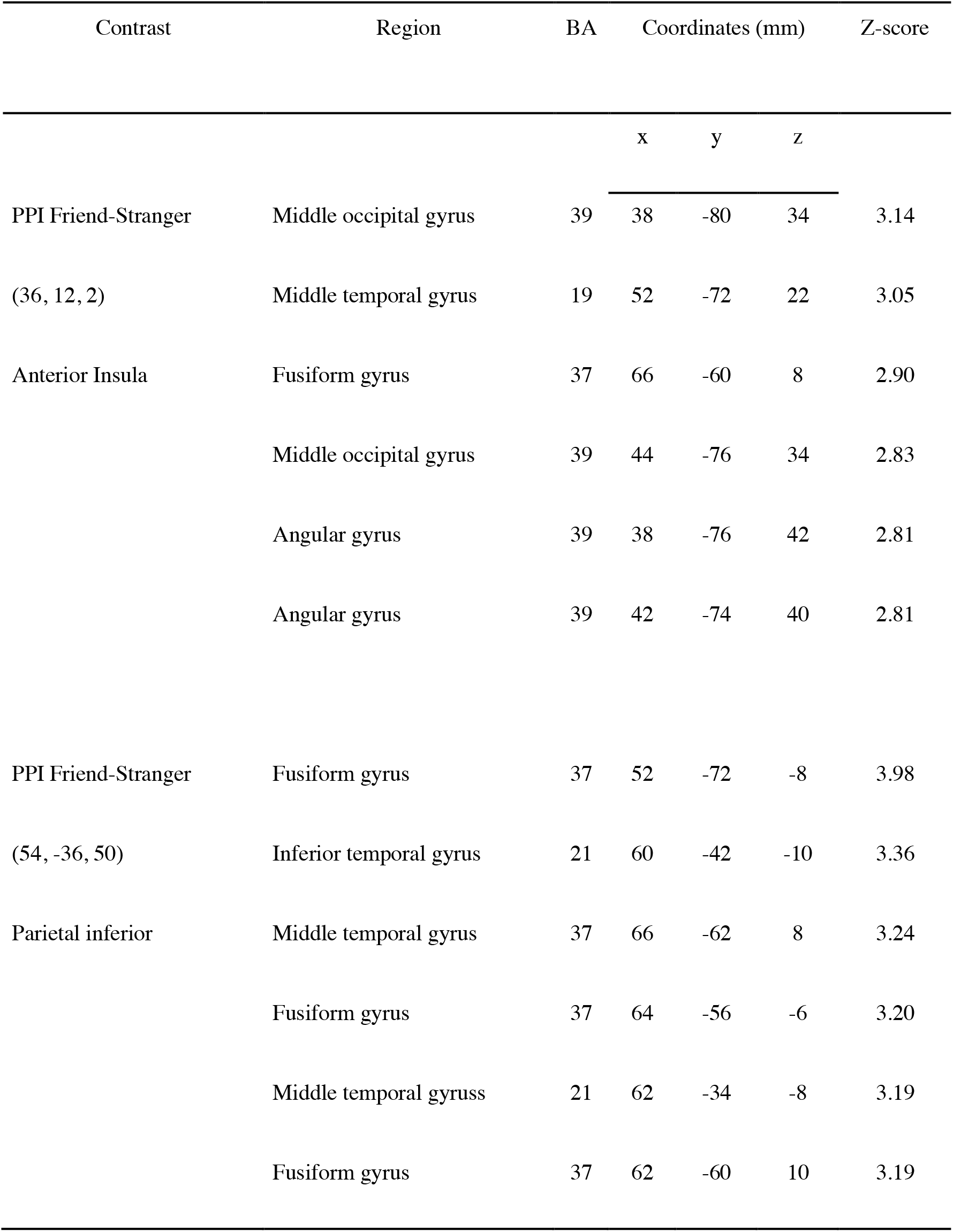
PPI results

## 4. Discussion

This study aimed to investigate the brain dynamics related to trust anticipation and social closeness. We found that subjects in the Trust Game decided to pay back more to Friends than to a Stranger and the Computer. During the anticipation of the Stranger’s decision, we found higher activity in the caudate nucleus and putamen than the Computer’s decision, while there was higher activity in the lingual gyrus, angular gyrus, cuneus, precuneus, putamen and anterior insula in the anticipation of the Friend’s decision vs the Computer. We also found higher activity in the supplementary motor area, middle frontal gyrus, parietal lobe and angular gyrus when anticipating the Friend’s decision vs the Stranger’s. Finally, with the psychophysiological interaction analysis we found higher functional connectivity between the right anterior insula and the posterior fusiform gyrus, when anticipating Friend’s decision vs the Stranger’s. Our results suggest that the anterior insula is sensitive to (dis)trust anticipation from a human peer, regardless of their social closeness, and that it interacts differentially with the middle occipital and temporal gyrus, as well as the fusiform gyrus, depending on the social closeness.

We propose that the activation pattern revealed by the Friend vs. Computer contrast, particularly the activity of the AIns, can be interpreted as the experience of internal conflict in the face of potential distrust of the closest partner. In a similar study, in which participants played the Dictator Game with close peers who varied in the relationship valence (friends vs. dislike peers), the authors found that people who were less prosocial toward their friends compared to their dislike peers, had greater activation of the Supplementary Motor Area (SMA) and AIns during the game (Schreuders et al., 2018). In this study, acting prosocial towards a friend is interpreted as a social norm, and not acting according to it induces internal conflict. We consider that the internal conflict could indicate both, acting against a social norm and norms deviation anticipation by other agents. Studies that indicate that being treated unfairly coincides with the AIns activity could support this idea (Güroglu et al., 2014; Sanfey et al., 2003). Likewise, we propose that the AIns activity that occurs even before the subjects can act reciprocally, is sensitive to social information. It is very highly possible that these social inputs, such as the partner’s closeness or the uncertainty regarding their behavior, have a significant impact on the process of assigning values to alternatives, that underlie reciprocal or selfish decision-making, and that it is probably occurring during the anticipation phase (Lim et al., 2013; Rangel & Hare, 2010). Thus, the greater AIns activity during the anticipation of a human truster it is possible explained through internal conflict, caused by the possibility of being treated unfairly by the partner with the greatest social closeness. It should be noted that despite the possible internal conflict, we found that the greater the truster’s social closeness, the greater were the reciprocal decisions of our subjects. Although this result could seem contrary to the internal conflict hypothesis, the neural dynamics between AIns and Fusiform gyrus, as well as their role in regulating the aversive experience through analysis of the underlying motivations, could explain these behavioral results.

The greater activation of the Fusiform Gyrus (FG) in trust anticipation from friends compared to strangers is an interesting result. The FG, which underlies our ability to process faces to interact in a socially appropriate way, has also been involved in the anticipation of monetary rewards (Dillon et al., 2008), and has been found to increase their response to emotional stimuli with low and high social complexity (Geday et al., 2003). It has been suggested that there is a neural network that includes the posterior FG and the inferior occipital gyrus, which specializes in identifying visual signals of high emotional importance. In addition, it has been proposed that the functioning of this network is fundamental during empathic reactions underlying a social interaction (Geday et al., 2003), and that its alterations, could partially explain the social dysfunction observed in patients with autism spectrum disorders (Pierce & Redcay, 2008). Thus, the differential activation of the FG during the anticipation of the high vs low closeness partner could suggest the occurrence of an attentional process, which, motivated by the uncertainty regarding the peers’ decisions, is probably more demanding during the anticipation of the friend compared to the stranger. Attention during anticipation would be essential when monitoring the decisions of the in-group partner because receiving their (dis)trust could be perceived as more rewarding (or punishing) than receiving the decision consequences of an out-group person.

Using a psychophysiological interaction (PPI) approach we detected greater functional connectivity between the right AIns and the middle temporal sulcus, during decision’s anticipation from an in-group versus an out-group trustor. Likewise, during the mentioned psychological context, the right AIns exhibited greater interaction with the right parietal cortex and the superior division of the lateral occipital cortex. These results suggest that: 1) The AIns is sensitive to (dis)trust anticipation from a human peer, regardless of their social closeness, presumably to encode aversive states caused by potential deviations from the trust social norm (Güroglu et al., 2014; Sanfey et al., 2003), and 2) the AIns interacts differentially, depending on the social closeness, especially with the middle temporal gyrus, possibly to regulate the aversive experience by analyzing the intentions and objectives underlying the partner’s trust (Geday et al., 2003).

The neural dynamics observed in our study may be understood through the Punishment Neuropsychological Framework (PNF) (Krueger & Hoffman, 2016). Although punishment was not directly evaluated in the present work, our experiment allowed us to assess the anticipation of behaviors that might warrant punishment. The PNF proposes that humans frequently punish others whose behavior deviates from the norms, e.g., direct victims of violations can retaliate against their aggressors (second party punishment), even people who were not directly harmed are willing to punish who transgresses (third party punishment) (Buckholtz & Marois, 2012). How willing people are to punish depends on the harm was done and the intentions of the offender (Treadway et al., 2014). Brain regions belonging to the Salience Network (SN) and the Default Mode Network (DMN) are thought to be involved in the detection of deviations from social norms, also encoding of the damage’s severity, and the intention assessment (Bressler & Menon, 2010; Krueger & Hoffman, 2016). In particular, it has been proposed that the AIns represents social norms and generates aversive experiences depending on whether there is a violation, or harm threat (Krueger et al., 2020), while the Temporal Parietal Junction (TPJ) and the temporal sulcus represent the nature of the social interaction, in terms of the intentions or objectives of the transgressor (Koldewyn & Kanwisher, 2018; Pelphrey et al., 2004; Schiopu, 2016; Shultz et al., 2011) and even encode social distance (Parkinson et al., 2017; Strombach et al., 2015). Our whole-brain results are congruent with the AIns’ role in the information encoding related to anticipation of deviations from social norms (Xiang et al., 2013), however, we also show that the AIns interacts with the temporal regions depending on the trustor’s social closeness. The functional coupling between these regions that we observed during trust anticipation from a high vs low social closeness partner, could reflect the information flow between the SN and the DMN (particularly between the AIns, the TPJ, and the posterior Superior Temporal Sulcus), necessary to modulate the differential negative affect, produced by the uncertainty regarding the friend’s behavior compared to the stranger. It is reasonable to speculate that the effect of the social cognition network when monitoring the intention of a proximate behavior, decreases the response of SN’s regions (e.g. the amygdala), in a similar way to high-level regulatory strategies, affect emotional experiences (Diekhof et al., 2011; Treadway et al., 2014). A related result to the previous idea was that the whole brain analysis detected basal ganglia activity during the anticipation of the human trustor’s decisions relative to the non-social control, however, this region was absent in our contrast to evaluate the difference between friend and stranger’s anticipation. In agreement with the PNF and the supposed modulation of affect, it is plausible that the basal ganglia response could have been inhibited, yet, this hypothesis could be evaluated in future studies.

### 4.1. Strengths and limitations

Our study has several strengths, beyond just focusing on whole brain activations during the anticipation of truster’s decisions, the PPI model we used allowed us to evaluate the functional interaction of this region with other brain structures. PPI allows investigating not only individual regions involved in the task but also how is the information flow between brain areas and how functional regions change their connectivity in different psychological contexts (Di et al., 2020; O’Reilly et al., 2012). The PPI analysis strengthens the study in terms of the sensitivity of our neuroimaging findings, while reducing the impact of our sample size, which could be considered relatively small for imaging studies. Another strength of the study was the participation of real-life partners (close friends), who although their decisions were programmed, the expectation of their presence during the game increases the validity of the task and results (Schreuders et al., 2018). However, we also had important limitations that must be considered. The PPI does not allow establishing the directionality of the information flow between brain areas (Fareri et al., 2020), so we do not know if the AIns and parietal inferior receive or send information from the areas they interact with. Although we tried not to make inferences regarding the direction of the relationship between regions, our hypothesis that the mentalization network modulates the aversive experience in the face of the potential distrust of the friend could assume directionality of the information from the temporal areas to AIns. Therefore, future studies should empirically evaluate this question through effective connectivity methods such as Granger causality analysis or dynamical causal modeling (Treadway et al., 2014). Another limitation of the present study was the use of hypothetical monetary rewards rather than real, it could be argued that the subjects might not be sufficiently motivated by the consequences of decisions in the game. However, in the research literature on decision making, there are numerous studies that have explicitly addressed the difference between hypothetical versus real monetary rewards, without finding effects of the type of reward in self-control, temporal or social discount tasks (Johnson & Bickel, 2006; Locey et al., 2011). Considering the aforementioned studies, as the findings of this work have theoretical congruence, there are few reasons to believe that other types of incentives would have led to different results.

## 5. Conclusions

Human social life success depends to a large extent on people trusting and taking risks together, with the purpose of achieving objectives that otherwise would fall short of reach. When it comes to interactions between members of the same group, the default is to anticipate the trust and reciprocity of close others like our friends. However, in societies as numerous and complex as human ones, it is frequent that many of our interactions occur with people hardly known or strangers. Although it may be less frequent, it is also possible that close group members prefer, in some circumstances, not to take the risk of placing their trust in us. In this way, anticipating the trust of another individual requires that we be able to selectively attend to socially relevant stimuli, such as closeness or other’s past decisions, to generate adequate expectations regarding their behavior. And in case of anticipating a deviation from a social norm, analyze the motivations or objectives of the involved person, and make a motivated prosocial or proself decision. This complicated neuropsychological process requires the information flow between neural regions sensitive to social nature data, such as the AIns and temporal gyrus, which functionally interact to signal the other person closeness and modulate the aversive response that occurs as a consequence of potential distrust.

## Acknowledgements

This study was possible thanks to the support of the Instituto Nacional de Psiquiatría Ramón de la Fuente Muñiz.

